# DeepSSV: detecting somatic small variants in paired tumor and normal sequencing data with convolutional neural network

**DOI:** 10.1101/555680

**Authors:** Jing Meng, Brandon Victor, Zhen He, Agus Salim

## Abstract

**Motivation:** It is of considerable interest to detect somatic mutations in paired tumor and normal sequencing data. A number of callers that are based on statistical or machine learning approaches have been developed to detect somatic small variants. However, they take into consideration only limited information about the reference and potential variant allele in both samples at a candidate somatic site. Also, they differ in how biological and technological noises are addressed. Hence, they are expected to produce divergent outputs.

**Results:** To overcome the drawbacks of existing somatic callers, we develop a deep learning-based tool called DeepSSV, which employs a convolutional neural network (CNN) model to learn increasingly abstract feature representations from the raw data in higher feature layers. DeepSSV creates a spatially-oriented representation of read alignments around the candidate somatic sites adapted for the convolutional architecture, which enables it to expand to effectively gather scattered evidences. Moreover, DeepSSV incorporates the mapping information of both reference-allele-supporting and variant-allele-supporting reads in the tumor and normal samples at a genomic site that are readily available in the pileup format file. Together, the CNN model can process the whole alignment information. Such representational richness allows the model to capture the dependencies in the sequence and identify context-based sequencing artifacts, and alleviates the need of post-call filters that heavily depend on prior knowledge. We fitted the model on ground truth somatic mutations, and did benchmarking experiments on simulated and real tumors. The benchmarking results demonstrate that DeepSSV outperforms its state-of-the-art competitors in overall F_1_ score.

**Availability and Implementation:** https://github.com/jingmeng-bioinformatics/DeepSSV

**Supplementary information:** Supplementary data are available at online.

## 1 Introduction

Due to advancements in high-throughput sequencing technologies, it is of considerable interest to detect somatic mutations from tumor and matched normal sequence data in the cancer research community (**??**). By comparing the sequences in tumor cells with those in normal cells from the same individual, a list of potential somatic mutations are identified and could be used by researchers for further validation studies to confirm their causal roles in tumor development. As subsequent validation experiments are expensive and time consuming, improving prediction accuracy of computational methods is essential (**?**). To achieve this goal, a major challenge is how to accurately model biological and technological noises, including intra-tumor heterogeneity, sample contamination, uncertainties in base sequencing and read alignment (**???**).

A number of callers have been developed to detect somatic single nucleotide variants (SNVs) and insertions and deletions (indels), which basically fall into two categories: statistical-based (**???????**) and machine learning-based approaches (**????**). MuTect2 (**?**) combines two separate but dependent Bayesian classifiers with local assembly of haplotypes to classify variants as somatic (no evidence of existence in normal) or germline (sufficient evidence of existence in normal). Possible false predictions are then removed by hard filters. EBCall (**?**) empirically evaluates sequencing errors from multiple non-paired normal samples with a Beta-binomial distribution, and includes it as prior information in a Bayesian model to discriminate somatic mutations from sequencing noise. Strelka (**?**) employs the strategy that considers the normal sample to be a mixture of germline variation and noise, and the tumor sample to be a mixture of the normal sample and somatic variation, which expands the range of detectable variant allele fractions (VAFs) without requiring purity estimation. Developed for the same purpose, somatic callers vary in the following ways: the diversity level of factors taken into consideration; in the way noises are estimated; in the threshold used to call a mutation; and in the stringency level of post-call filters to exclude potential false positives. As a result, it is not surprising to see low agreement between these callers (**?????**).

MutationSeq (**?**) is a tool that implements standard machine learning algorithms to detect somatic SNVs, which is followed by SomaticSeq (**?**), SNooPer (**?**) and ISOWN (**?**). The features used to train the classifiers on a set of ground truth somatic positions are either extracted from packages such as GATK (**?**) and Samtools (**?**) or defined by developers themselves to boost weak signals in the tumor and identify systematic errors. Such classifiers are flexible in that they avoid a need for strong assumptions about underlying mechanisms and their performance can be improved by adding hand-crafted features, but it is a labor-intensive process and requires domain knowledge to derive discriminative features.

Deep learning automates this critical step of feature selection and learns increasingly abstract feature representations from the raw data in higher feature layers by using deep architectures (**???**). Given the advantages that deep networks can operate on the sequence directly without requiring pre-defined features, deep leaning approaches have been successfully applied in bioinformatics (**???**). However, in the area of variant detection, we are only aware of two deep learning-based approaches. The first is Deepvariant (**?**) that uses a deep convolutional neural network (CNN) to call germline SNPs and small indels in normal genomes. Deepvariant encodes mapped reads at each candidate genomic site in normal and reference data as an image, and feeds it to a CNN model pre-trained for image classification tasks to calculate the genotype likelihoods. It performs well on germline variant calling for different sequencing technologies. Nevertheless, it is sub-optimal to transform alignment information into images as the four-channel images cannot effectively represent all of the available information. The other work is Kittyhawk (**?**) for detection of somatic SNVs with ultra-low VAFs in tumor liquid biopsies such as cell-free circulating DNA (cfDNA).

The predictive models that are based on statistical or machine learning approaches in previous works account for only limited information (the VAF, the depth, the base and mapping qualities) about the reference and potential variant allele in both samples at a candidate genomic site. The disadvantages of such models are twofold. First, evidences about a genomic site being somatic are not confined to the site but scattered apart. However, existing models do not scale to incorporate spatially-oriented evidences, which leaves systematic errors unidentified (**?**). Second, regarding the alignment information at a site, these models can only see a part of it instead of a whole picture. Together, it is not enough to do robust inference. The only somatic caller based on deep learning approach, Kittyhawk (**?**) considers only the genomic sites with a copy of non-reference variant allele regardless of the depth, and encodes the mapping information of the supporting read to train a CNN classifier. This read-centric approach cannot be applied to identify somatic mutations with multiple variant-allele-supporting reads to fully understand the temporal order of somatic events and tumor clonal architecture.

To address the disadvantages of existing approaches, we develop a tool called DeepSSV to call somatic SNVs and small (1-50 bp) indels with more than one supporting read, which uses the CNN model to discriminate signal from noise. Our contributions are as follows: (1) we address the first disadvantage by creating a spatial representation of mapping information around a candidate somatic site, which allows the solution to effectively gather scattered evidences about a site being somatic; (2) we encode the mapping information of both reference-allele-supporting and variant-allele-supporting reads in the tumor and normal samples that are readily available in the pileup format file, so that the CNN model can see the alignment information of every read. Such representational richness enables the capture of dependencies in the sequence and detection of context-based sequencing artifacts, and alleviates the need to design post-call hard filters that are heavily dependent on prior knowledge. We trained the model on experimentally-validated somatic mutations, and ran benchmarking experiments on simulated and real tumors. The results show that DeepSSV outperforms state-of-the-art somatic callers in overall F_1_ score.

## 2 Methods

### 2.1 Workflow of DeepSSV

We show the workflow of DeepSSV for somatic small variant prediction in Figure 1. DeepSSV takes as input a mixed pileup file generated by samtools from tumor and normal BAM files. Supplementary Material shows how to convert these two BAM files to a mixed pileup file required by DeepSSV. DeepSSV first operates on each genomic site independently to identify candidate somatic sites. Next it encodes the mapping information around the candidate somatic sites into an array. Each array is a spatial representation of mapping information adapted for the convolutional architecture. Then the CNN model trained on experimentally-validated somatic events evaluates the information in these arrays to obtain additional support for true positives and filter false positive predictions. Finally, potential somatic small variants determined by the CNN model are generated in the variant call format (VCF). The following is a detailed breakdown of how DeepSSV works.

**Fig. 1.**
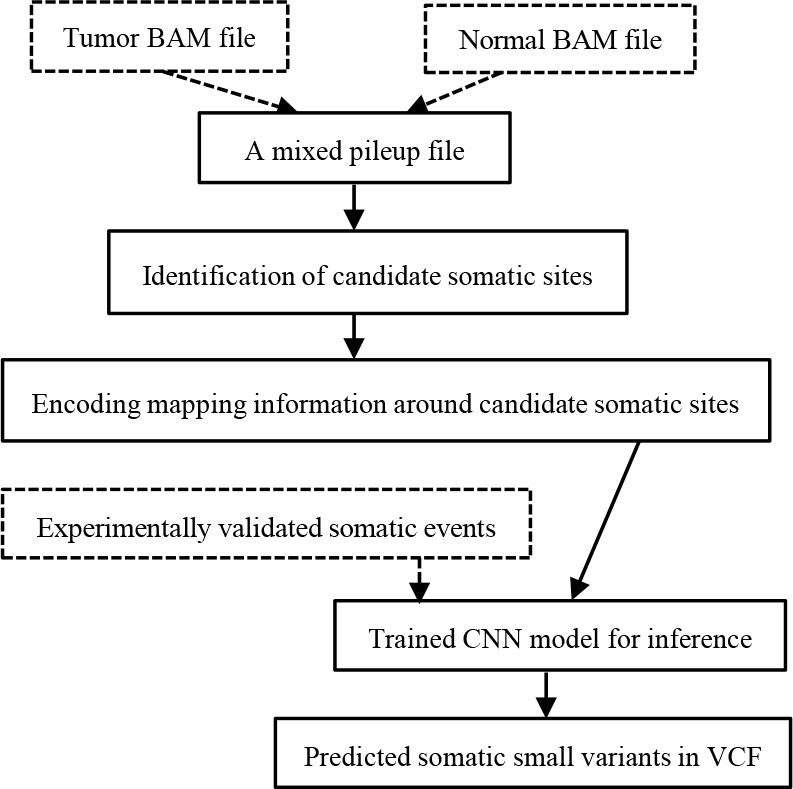
Workflow of DeepSSV. Outside DeepSSV’s workflow, tumor and normal BAM files are transformed into a mixed pileup file by samtools, and the CNN model is trained on experimentally-validated somatic events. These two steps are represented using dashed lines. DeepSSV first scans each genomic site to identify candidate somatic sites, then incorporates the mapping information around each candidate somatic site into an array, which is provided to the trained CNN model for inference.

#### 2.1.1 Identification of candidate somatic variants

The first step of DeepSSV’s workflow is identifying candidate somatic small variants for further evaluation by the CNN model. Each position in the reference genome is considered independently. For all reads overlapping a genomic site, we include only the following reads when counting the number of occurrences of each type of allele: reads with reference base, non-reference base, an insertion with a specific sequence and a deletion with a specific length. If there are multiple non-reference alleles in the tumor at a genomic site, we choose the top one in terms of allelic abundance. When converting BAM file to pileup format file by Samtools (**?**), we exclude low quality reads whose mapping quality and base quality at a specific site are low (default *≤* 10 and *≤* 13 respectively) (**?**), and any unusable reads such as duplicates, secondary alignments and improper pairs for paired-end sequencing.

After filtering away low quality alignments, we focus on each genomic site to determine its possible state. The criteria used to choose candidate somatic sites are: (1) the depths in both the normal and tumor are equal to or larger than 10; (2) the base in the reference genome is standard (ACGT); (3) the VAF of a variant allele in the tumor is larger than a threshold (default *≥* 0.075), and is larger than that in normal if the variant allele exists in normal; (4) the copies of a variant allele in the tumor should be present in both the forward and reverse strand, because extreme strand bias induced by polymerase chain reaction (PCR) duplicates from the sequencing machines is an indicator of potential false positive (**?**); (5) there is a low number of occurrences of a variant allele and a low VAF in the normal (default < 2 and < 0.03 respectively) (**?**); and (6) the length of the inserted or deleted sequence in the tumor is smaller than 50. We limit the size of somatic indels to match the ground truth somatic indels that we used to train the CNN model.

#### 2.1.2 Encoding mapping information around candidate somatic sites

The second step of DeepSSV is incorporating mapping information around candidate somatic sites and encoding such information into an array to feed into the CNN model. Supplementary Figure S1 illustrates how to carry out the incorporation of mapping information into an array. In order to fully capture the mapping information and systematic artifacts in next-generation sequencing (NGS) data, we generate an array for each candidate somatic site to represent spatially-oriented read alignment. We encode DNA nucleotides by one-hot encoding: A = (0 0 0 0 1), T = (0 0 0 1 0), G = (0 0 1 0 0), C = (0 1 0 0 0), N = (1 0 0 0 0). The array for each candidate somatic site has 221 columns centered on the candidate site, as one column represents one flanking genomic site to the left or right of the candidate site. The number of columns can be changed by users. In each column, the corresponding reference base in the human genome are put in the first 5 rows. Instead of transforming the alignments to allelic accounts, we supply mapping information of each individual base covering the genomic site to the CNN model. For each covering base, its mapping information is readily available in the pileup format file and takes 14 rows in total. The mapping information contained in each array is read base, strand information (forward or reverse strand), CIGAR string, mapping quality, base quality and distance to the start of the mapping read. The CIGAR string or alignment string describes how the read base is aligned to the reference. In our work only four CIGAR operations are used, which are sequence match (=), sequence mismatch (X), insertion to the reference (I) and deletion from the reference (D). We use one-hot encoding to represent the categorical features as well. Although the bases for the deletion come from the reference genome, we do not include them at the corresponding sites as the qualities for them are fake and they do not actually exist in the called read. Also, inserted bases are between reference sites, so they do not appear in the array. As data normalization is able to optimize model performance (**?**), to scale the values to be in the interval [0, 1], we divide base distance to the start of the mapping read by the read length, and use the base-calling error probability rather than Phred quality score to indicate uncertainties in base calling and read alignment. To maintain the same length of each column, if there are less mapped reads than default (100) at the corresponding site, we apply a padding with a value that is not a real input so that it does not have an impact on the model’s behavior. We append the mapping information of reads in normal to that in tumor. In total, there are 2805 rows in one column.

#### 2.1.3 The CNN model

The array for a candidate somatic site is two-dimensional (2D) and contains raw mapping information instead of hand-crafted features, which poses a challenge for classical machine learning algorithms. Given this consideration, we turn to the CNN to learn increasingly abstract feature representation from the raw data. A CNN is typically composed of convolutional layers and pooling layers, which are followed by fully connected layers. It performs convolutional computation on small regions of the output from the previous layer and shares parameters between regions of a feature map (**?**). Compared with a purely fully-connected model, the CNN architecture leads to a smaller number of model parameters.

We use the CNN model implementation to detect somatic mutations as it allows direct operation on the mapping information of reads. The architecture of our CNN model is shown in Figure 2. The array fed into the CNN model has 2805 rows and 221 columns. Since there is no spatial representation of the mapping information of reads at a site in each column of an array, we apply 1D convolutional operations in the convolutional layers. Tri-nucleotide context (*±*1 bp) at a genomic site contains information about mutational signatures and underlying mutational processes (**?**). To gather equal information from both directions around a genomic site, we use kernel/filter size of 3 and stride of 1 for the convolutional operation. After two successive convolutional layers, to keep the most important mapping information in small spatial genomic regions, we do down-sampling by max-pooling with kernel/filter size of 2 and stride of 2.

**Fig. 2.**
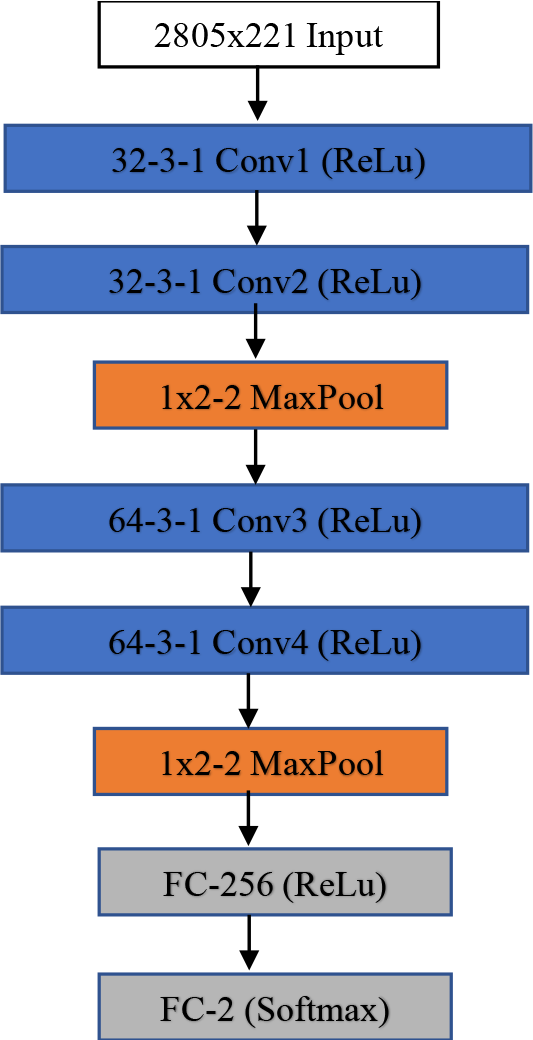
The architecture of our CNN model. The input corresponding to each candidate somatic site has 2805 rows and 221 columns. ‘32-3-1 Conv1’ denotes the first 1D convolutional layer with 32 filters of size 3 and stride 1. ‘1×2-2 MaxPool’ denotes a max pooling layer implementing down-sampling over 1×2 regions with stride 2. ‘FC-256’ and ‘FC-2’ denote a fully connected linear layer with 256 and 2 neurons respectively. 2 is the number of classes that the model outputs. ‘ReLu’ and ‘Softmax’ denote different activation functions.

### 2.2 Datasets

#### 2.2.1 Real data

We used two real paired tumor and normal whole genome sequencing (WGS) datasets in our study. The first dataset is from a paired COLO829 and COLO829BL cell lines, which were sequenced by separate institutions with different library preparations to create a somatic reference (**?**). We downloaded the BAM files provided by Translational Genomics Research Institute (TGen) and downsampled the reads from ~80× to ~50× for both tumor and normal samples. The second dataset is corresponding to a case of medulloblastoma (MB) from the International Cancer Genome Consortium (ICGC) (**?**). The original FASTQ files are of diverse sequencing libraries as well. Library L.A gives relatively high evenness of coverage genome-wide, so we chose to use its sequencing files. Sequencing depths for tumor and normal samples are ~40× and ~30× respectively. The BAM files provided were not used directly. Instead, we first used BWA to map the reads in the downloaded FASTQ files to the human reference genome hg38/GRCh38 with default settings (**?**), then marked PCR and optical duplicates with Picard, and finally realigned the raw gapped alignment and adjusted base quality scores with GATK (**?**).

Most of the currently available ground truths of individual tumors are of small size, containing only hundreds to thousands of somatic events, which are not enough to train CNN models. Although transfer learning is an effective way of eliminating the requirement for large training examples (**?**), there are no appropriate pre-trained models that can be reused for our study, and it is very likely that fine-tuned models on small-sized balanced training data will produce a high number of false positives as non-somatic sites significantly outnumber somatic sites in tumor genomes. To overcome these drawbacks, one way is to create a large imbalanced training set which consists of limited validated somatic sites and overrepresented non-somatic sites. Training a model on imbalanced data is possible, but highly-skewed class distribution leads to compromised models’ performance (**?**). The COLO829 genome has a high mutational load (**?**). The current version of its ground truth genotype calls includes approximately 35,000 somatic SNVs and 400 somatic small indels, which enables us to create a large balanced training set to filter false positive predictions. On the contrary, the MB genome is lowly mutated, carrying only approximately one mutation per megabase (**?**), which serves as a somatic standard to evaluate models’ ability of distilling the signal from the noise. We lifted over the ground truth VCF files corresponding to these two genomes to GRCh38 to match the BAM files.

#### 2.2.2 Simulated data

Three simulated tumor genomes were employed for benchmarking as well (**?**). They were created by introducing somatic mutations into homozygous reference allele sites of the NA12878 genome that was well characterized to generate single nucleotide polymorphism (SNP) and homozygous reference genotype calls of high confidence. The paired pre-tumor or normal genomes differ in the library design and sample preparation, and the resulting simulated tumors differ in the mutation frequency across the genome, the number of sub-clones and the VAFs. Supplementary Tables S1 and S2 give detailed description of these paired normal and simulated tumor genomes. One of the simulated tumors has a mutation load for somatic SNVs of 5 mutations per megabase, and three cell sub-populations that correspond to expected VAFs of 0.2, 0.35 and 0.5. The other two simulated tumors have a mutation load for somatic SNVs of 10 mutations per megabase, and four cell sub-populations corresponding to expected VAFs of 0.1, 0.2, 0.35 and 0.5. The mutation load for somatic small indels of each simulated tumor is 10% of that of somatic SNVs. Within a tumor, each sub-population is in the same proportion. The performance of our classifier was estimated against these intact highly confident germline and reference sites, and simulated somatic sites in the NA12878 genome.

### 2.3 Experimental design

We divided the ground truth somatic sites of the COLO829 genome into training, validation and testing sets. We fitted the CNN model on the training set, optimized hyper-parameters on the validation set and evaluated the model’s performance on the test set. Since the complete set of validated somatic sites of the COLO829 genome is not currently available, we first ran MuTect2 (**?**) and Strelka2 (**?**) on the COLO829 genome, and considered the sites that failed the post-call filters of both of the two tools as non-somatic sites which were then used as part of the large balanced training set.

To estimate the generalization performance of the trained CNN model, we applied it to the MB genome whose sequencing and subsequent mapping error profiles are different from the training set, and compared its performance with four state-of-the-art somatic callers, Strelka2, Lancet (**?**), MuTect2 and Deepvariant. Furthermore, we compared the performance of our trained model and three somatic callers (Strelka2, Lancet and Deepvariant) on three simulated tumor genomes. When benchmarking with simulated tumors, we did not choose MuTect2 as no somatic sites passed its post-call filters, and focused on just the regions with highly confident genotype calls (**?**). Supplementary Material gives information about the parameters we used when running these somatic callers.

We calculated two statistics for benchmarking somatic callers, precision and recall. Precision is the fraction of the predicted somatic sites that are true positives. Recall is the fraction of ground truth somatic sites that are correctly identified. F_1_ score is the harmonic average of precision and recall, which describes the overall performance of a somatic caller. The definition of F_1_ score is:

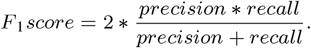

## 3 Results

### 3.1 Performance of DeepSSV on the COLO829 genome

We first examined the performance of our CNN model on the test set of the COLO829 genome. The test set contains 6846 ground truth somatic sites, 6769 of which are somatic SNVs. We used the default threshold to call a somatic site. Table 1 shows our model’s performance on the test set. Our model achieves comparable precision (0.9830) and recall (0.9791), and an overall F_1_ score of 0.9810. There are significant differences in the performance between somatic SNVs and somatic indels. The F_1_ score for somatic SNVs is about 0.36 higher than that for somatic indels. The AUC values for somatic SNVs and somatic indels are 0.9984 and 0.7830 respectively (Supplementary Figure S2). The differences are reasonable, considering that in the training set somatic SNVs greatly outnumber somatic indels (less than 300 sites).

**Table 1.** Accuracy metrics of DeepSSV on the COLO829 genome.

| Mutation type | # of calls | True Positives | Precision | Recall | F<sub>1</sub> score |
| --- | --- | --- | --- | --- | --- |
| Total | 6819 | 6703 | 0.9830 | 0.9791 | 0.9810 |
| Somatic SNVs | 6767 | 6663 | 0.9846 | 0.9843 | 0.9844 |
| Somatic indels | 52 | 40 | 0.7692 | 0.5195 | 0.6202 |

### 3.2 Benchmarking results on simulated tumor genomes

Next we ran benchmarking experiments on three simulated tumor genomes to investigate the performance of DeepSSV and three state-of-the-art somatic callers. Tables **??**-**??** and Supplementary Figures S3-S8 show the benchmarking results of these somatic callers. As is the case with the test set of the COLO829 genome, the performance of each somatic caller on somatic indels is worse than that on somatic SNVs. The highest F_1_ score (0.7463) on somatic indels is produced by DeepSSV on the simulated_tumor_1 genome (Table 2), while the highest F_1_ score on somatic SNVs is 0.9415, which is from Strelka2 on the simulated_tumor_2 genome (Table 3). The performance difference between the detection of somatic indels and somatic SNVs is the most obvious in Deepvariant. For instance, it detects 86.79% of simulated somatic indels, which is accompanied by a very low precision value of 0.0914, resulting in F_1_ score of 0.1654 (Table 3). In contrast, its F_1_ score on somatic SNVs of the same simulated tumor genome is 0.6375.

**Table 2.** Accuracy metrics of somatic callers on the simulated_tumor_1.

| Caller | # of calls | TP | Precision<br>Somatic indels | Recall | F<sub>1</sub> score | # of calls | TP | Precision<br>Somatic SNVs | Recall | F<sub>1</sub> score | Precision<br>Total | Recall | F<sub>1</sub> score |
| --- | --- | --- | --- | --- | --- | --- | --- | --- | --- | --- | --- | --- | --- |
| DeepSSV | 1331 | 919 | 0.6905 | 0.8118 | **0.7463** | 16411 | 14981 | 0.9129 | 0.9303 | **0.9215** | 0.8962 | 0.9225 | **0.9092** |
| Lancet | 1897 | 997 | 0.5256 | 0.8807 | 0.6583 | 16612 | 13817 | 0.8317 | 0.8580 | 0.8446 | 0.8009 | 0.8600 | 0.8294 |
| Strelka2 | 1897 | 1059 | 0.5582 | 0.9355 | 0.6992 | 16580 | 14854 | 0.8959 | 0.9057 | 0.9008 | 0.8617 | 0.9238 | 0.8916 |
| Deepvariant | 1663 | 835 | 0.5021 | 0.7376 | 0.5975 | 12751 | 10693 | 0.8598 | 0.6808 | 0.7599 | 0.7568 | 0.6870 | 0.7202 |

TP: True Positives.

**Table 3.** Accuracy metrics of somatic callers on the simulated_tumor_2.

| Caller | # of calls | TP | Precision<br>Somatic indels | Recall | F<sub>1</sub> score | # of calls | TP | Precision<br>Somatic SNVs | Recall | F<sub>1</sub> score | Precision<br>Total | Recall | F<sub>1</sub> score |
| --- | --- | --- | --- | --- | --- | --- | --- | --- | --- | --- | --- | --- | --- |
| DeepSSV | 953 | 494 | 0.5184 | 0.8821 | **0.6531** | 8759 | 8256 | 0.9426 | 0.9889 | **0.9652** | 0.9009 | 0.9822 | **0.9398** |
| Lancet | 907 | 495 | 0.5457 | 0.8839 | 0.6748 | 8455 | 7377 | 0.8725 | 0.8836 | 0.8780 | 0.8408 | 0.8836 | 0.8616 |
| Strelka2 | 927 | 530 | 0.5717 | 0.9464 | 0.7128 | 8833 | 8259 | 0.9350 | 0.9892 | 0.9613 | 0.9005 | 0.9865 | 0.9415 |
| Deepvariant | 5427 | 496 | 0.0914 | 0.8679 | 0.1654 | 10217 | 7302 | 0.7147 | 0.8746 | 0.7866 | 0.5003 | 0.8785 | 0.6375 |

TP: True Positives.

Overall, DeepSSV is the best-performing somatic caller in terms of F_1_ score when taking somatic indels and somatic SNVs as a whole. It yields the F_1_ score of 0.9092, 0.9398 and 0.9300 on the simulated_tumor_1, simulated_tumor_2 and simulated_tumor_3 genomes, respectively. Strelka2 is comparable to DeepSSV on the simulated_tumor_2 genome, but its overall F_1_ score decreases to 0.9077 on the simulated_tumor_3 genome (Table 4). When it comes to individual types of somatic mutations, DeepSSV outperforms its competitors on all of simulated tumor genomes when detecting somatic SNVs, but it lags behind Lancet and Strelka2 on the simulated_tumor_2 genome, and Strelka2 on the simulated_tumor_3 genome respectively when detecting somatic indels. The gaps in precision between DeepSSV and two somatic callers (Lancet and Strelka2) increase with recall after recall gets to about 0.8 (Supplementary Figures S5 and S7). However, the gaps are overestimated as the maximum recall values that Lancet and Strelka2 can achieve are about 0.9. Contrary to other somatic callers, precision of Strelka2 increases with recall before recall gets to about 0.6 (Supplementary Figures S3, S4 and S6).

**Table 4.** Accuracy metrics of somatic callers on the simulated_tumor_3.

| Caller | # of calls | TP | Precision<br>Somatic indels | Recall | F<sub>1</sub> score | # of calls | TP | Precision<br>Somatic SNVs | Recall | F<sub>1</sub> score | Precision<br>Total | Recall | F<sub>1</sub> score |
| --- | --- | --- | --- | --- | --- | --- | --- | --- | --- | --- | --- | --- | --- |
| DeepSSV | 1369 | 815 | 0.5953 | 0.7975 | **0.6817** | 14932 | 14421 | 0.9658 | 0.9338 | **0.9495** | 0.9347 | 0.9254 | **0.9300** |
| Lancet | 1627 | 894 | 0.5494 | 0.8747 | 0.6748 | 14965 | 12911 | 0.8627 | 0.8360 | 0.8491 | 0.8320 | 0.8384 | 0.8352 |
| Strelka2 | 1578 | 927 | 0.5874 | 0.9070 | 0.7130 | 16152 | 14593 | 0.9034 | 0.9449 | 0.9236 | 0.8754 | 0.9426 | 0.9077 |
| Deepvariant | 5878 | 791 | 0.1346 | 0.7740 | 0.2293 | 14181 | 10926 | 0.7705 | 0.7075 | 0.7376 | 0.5857 | 0.7135 | 0.6433 |

TP: True Positives.

### 3.3 Benchmarking results on the MB genome

Finally, we applied DeepSSV and four state-of-the-art somatic callers to the MB genome. Table **??** and Supplementary Figures S9 and S10 describe the benchmarking results of these somatic callers. Compared with simulated tumor genomes, a much larger number of false positive somatic small variants contaminate the prediction set of these somatic callers. Two factors contribute to the lower precision values. The first is that, as highly confident genotype calls are not available in difficult regions, the regions of the NA12878 genome where we introduced simulated somatic small variants are relatively easily accessible by sequencers and mappers (**??**). The second factor is that the MB genome has a low mutational load, which leads to a highly-skewed class distribution.

Again, DeepSSV leads in the overall F_1_ score (0.7229, Table 5). In sharp contrast, 90.20% of ground truth somatic small variants are called by Deepvariant, which comes at the expense of an extremely low precision value (0.0070). Three other competitors are tuned to achieve high sensitivity with compromising precision. The overall recall values for MuTect2, Lancet and Strelka2 are 0.9528, 0.8469 and 0.9668 respectively. Although post-call filters corresponding to them are designed with the aim to exclude potential false positives, undesired precision values are yielded (0.1684 for MuTect2, 0.6123 for Lancet and 0.3795 for Strelka2). Without post-call filters available, the PRROC curve of DeepSSV on somatic SNVs is similar to its three competitors, MuTect2, Lancet and Strelka2 (Supplementary Figure S10). As on the simulated_tumor_2 genome, Lancet and Strelka2 have higher AUC values for somatic indels than DeepSSV (Supplementary Figure S9). Considering the limitations in sequencing and mapping technologies and the somatic callers used to identify ground truth somatic sites, it is impossible to think that the ground truth set really displays all somatic small variants (**?**). Hence, it is very likely that the precision values of somatic callers are underestimated.

**Table 5.** Accuracy metrics of somatic callers on the MB genome.

| Caller | # of calls | TP | Precision<br>Somatic indels | Recall | F<sub>1</sub> score | # of calls | TP | Precision<br>Somatic SNVs | Recall | F<sub>1</sub> score | Precision<br>Total | Recall | F<sub>1</sub> score |
| --- | --- | --- | --- | --- | --- | --- | --- | --- | --- | --- | --- | --- | --- |
| DeepSSV | 170 | 98 | 0.5765 | 0.3828 | **0.4601** | 1033 | 750 | 0.7261 | 0.8456 | **0.7813** | 0.7049 | 0.7419 | **0.7229** |
| MuTect2 | 2853 | 234 | 0.0820 | 0.9141 | 0.1505 | 3612 | 855 | 0.2367 | 0.9639 | 0.3800 | 0.1684 | 0.9528 | 0.2862 |
| Lancet | 395 | 192 | 0.4860 | 0.75 | 0.5898 | 1186 | 776 | 0.6543 | 0.8748 | 0.7486 | 0.6123 | 0.8469 | 0.7107 |
| Strelka2 | 1100 | 247 | 0.2245 | 0.9648 | 0.3642 | 1812 | 858 | 0.4735 | 0.9673 | 0.6357 | 0.3795 | 0.9668 | 0.5451 |
| Deepvariant | 96551 | 248 | 0.0026 | 0.9688 | 0.0052 | 58192 | 783 | 0.0135 | 0.8828 | 0.0266 | 0.0070 | 0.9020 | 0.0139 |

TP: True Positives.

## 4 Conclusion

Accurate identification of somatic mutations has the potential to improve downstream cancer diagnosis and treatment. A number of callers that are based on statistical or machine learning approaches have been developed to detect somatic small variants from tumor and matched normal sequence data. These callers differ in how biological and technological noises are addressed, and in the stringency level of post-call filters to exclude potential false positives. Therefore, it is not surprising that the predictions tend to have low concordance. Moreover, the existing somatic callers take into consideration only limited information (the VAF, the depth, the base and mapping qualities) about the reference and potential variant allele in both samples at a candidate somatic site.

Deep learning automates the feature engineering process. It uses deep architectures to learn increasingly abstract feature representations from the raw data in higher feature layers. Given the advantages that deep networks can operate on the sequence directly without requiring pre-defined features, we have proposed a deep learning-based tool called DeepSSV, which employs the CNN model to discriminate somatic mutations from biological and technological noises. DeepSSV gathers scattered evidences by creating a spatial representation of mapping information around candidate somatic sites that are readily available in the pileup format file. Moreover, DeepSSV encodes the mapping information of both reference-allele-supporting and variant-allele-supporting reads in the tumor and normal samples at a genomic site, so that the CNN model can see the whole mapping information.

We used the ground truth genotype calls of the COLO829 genome to create a large balanced training set, and fitted the CNN model on the training set. Then we assessed the model’s performance on the test set, which comes from the same genome as the training set. To investigate the generalization ability of our CNN model, we did benchmarking experiments on three simulated tumors and a lowly mutated real tumor. The benchmarking results show that DeepSSV, which does not have post-call filters, outperforms state-of-the-art somatic callers in overall F_1_ score.

## Acknowledgements

We would like to thank National Computational Infrastructure of Australia for its High Performance Computing (HPC) Systems and Cloud Computing Systems.

