## supplementary data for "DeepSSV: detecting somatic small variants in paired tumor and normal sequencing data with convolutional neural network"

*Corresponding author

**Converting tumor and normal BAM files to a mixed pileup file**

DeepSSV takes as input a mixed pileup file generated by samtools from tumor and normal BAM files. The following command shows how to convert these two BAM files to a mixed pileup file required by DeepSSV:

samtools mpileup -B -d 100 -f /path/to/ref.fasta [-l] [-r] -q 10 -O -s -a /path/to/tumor.bam /path/to/normal.bam > /path/to/mixed_pileup_file

For the case of applying DeepSSV on a part of the whole genome, increase the BED entry by 110 base pairs in each direction, and specify the genomic region via the option -l or -r.

**Parameters of somatic callers**

MuTect2 (version 2.1): We first ran MuTect2 without dbSNP and COSMIC files, and then ran FilterMutectCalls to label false positives with a list of failed filters and true positives with PASS. The records with ‘PASS’ in the FILTER field were used for analysis.

Lancet (version 1.0.5): We ran Lancet with default parameters. The records with ‘PASS’ in the FILTER field were used for analysis.

Strelka2 (version 2.9.2): Strelka2 was run with default settings. We used the predictions in the VCF files of somatic indels and somatic SNVs that pass the post-call filters for analysis.

Deepvariant (model version 0.6.0): Deepvariant was run on the tumor and matched normal BAM files separately. We considered to be somatic sites the predictions that are present in the VCF file from the tumor genome but not in the VCF file from the matched normal genome, and used these somatic sites for analysis.

**Supplementary Figures**

Reference

base

Read base

**Genomic context**

Strand information

CIGAR string

Mapping quality

Base quality

Base position on reads

Flanking sites

**Supplementary Figure S1.** Incorporating mapping information around candidate somatic sites into an array to feed into the convolutional neural network (CNN) model. The array corresponding to a candidate somatic site has 2805 rows and 221 columns centered on the candidate site. In each column, the corresponding reference base in the human genome and the mapping information (read base, strand information, CIGAR string, mapping quality, base quality and distance to the start of the mapping read) of covering reads take 2805 rows in total.

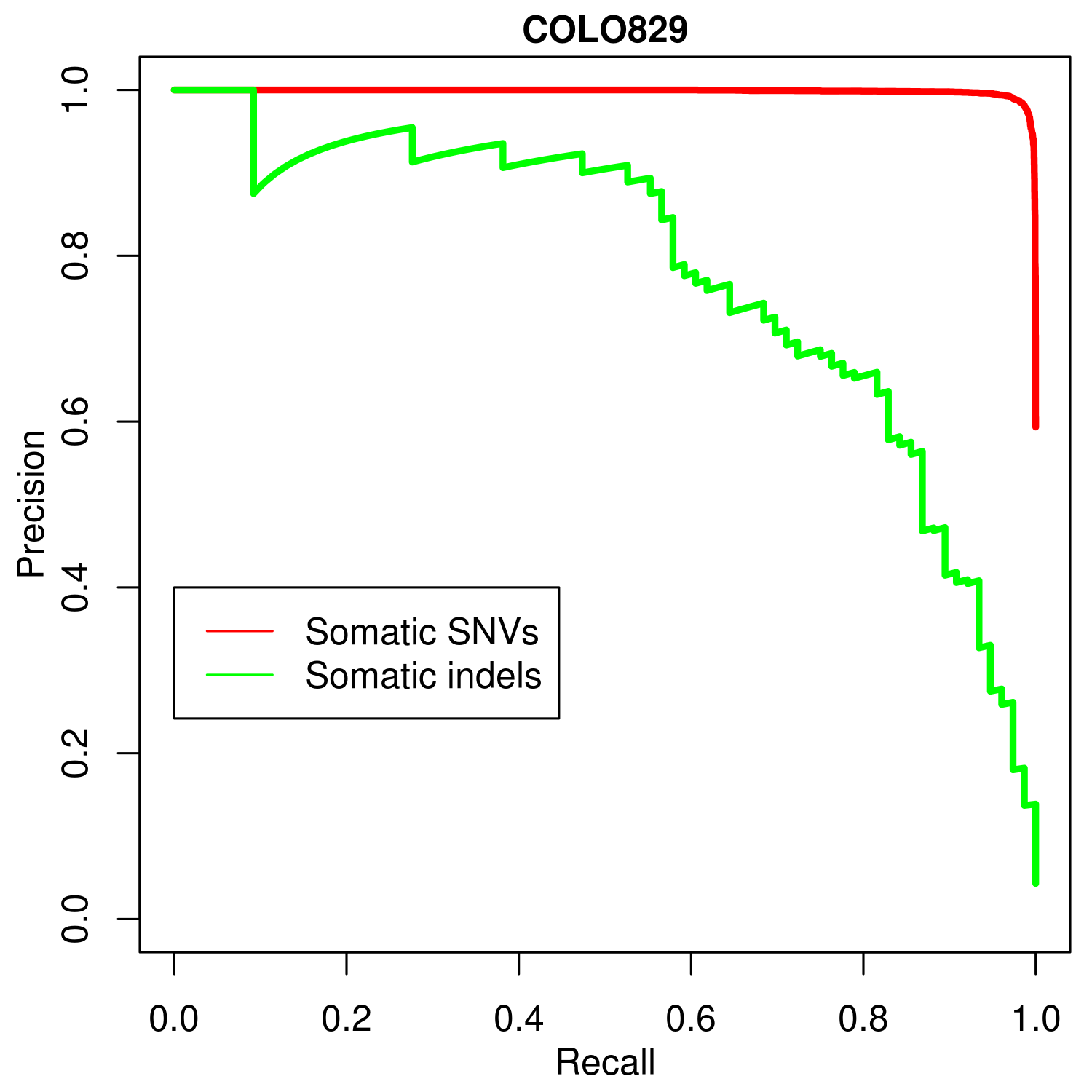

**Supplementary Figure S2.** PRROC curve of DeepSSV on the test set of the COLO829 genome. Red and green lines correspond to somatic SNVs and somatic indels, respectively.

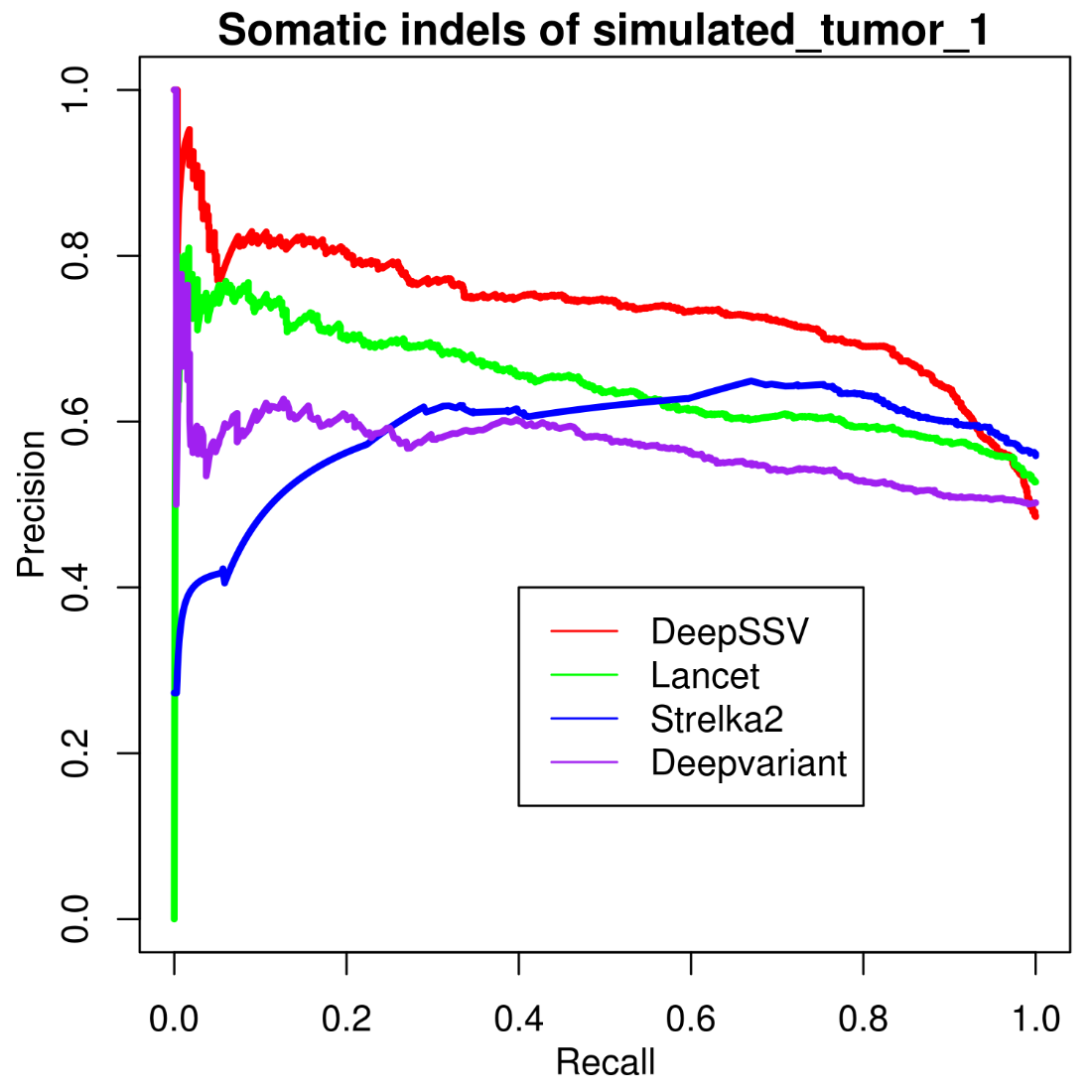

**Supplementary Figure S3.** PRROC curve of somatic callers on somatic indels of the simulated_tumor_1 genome.

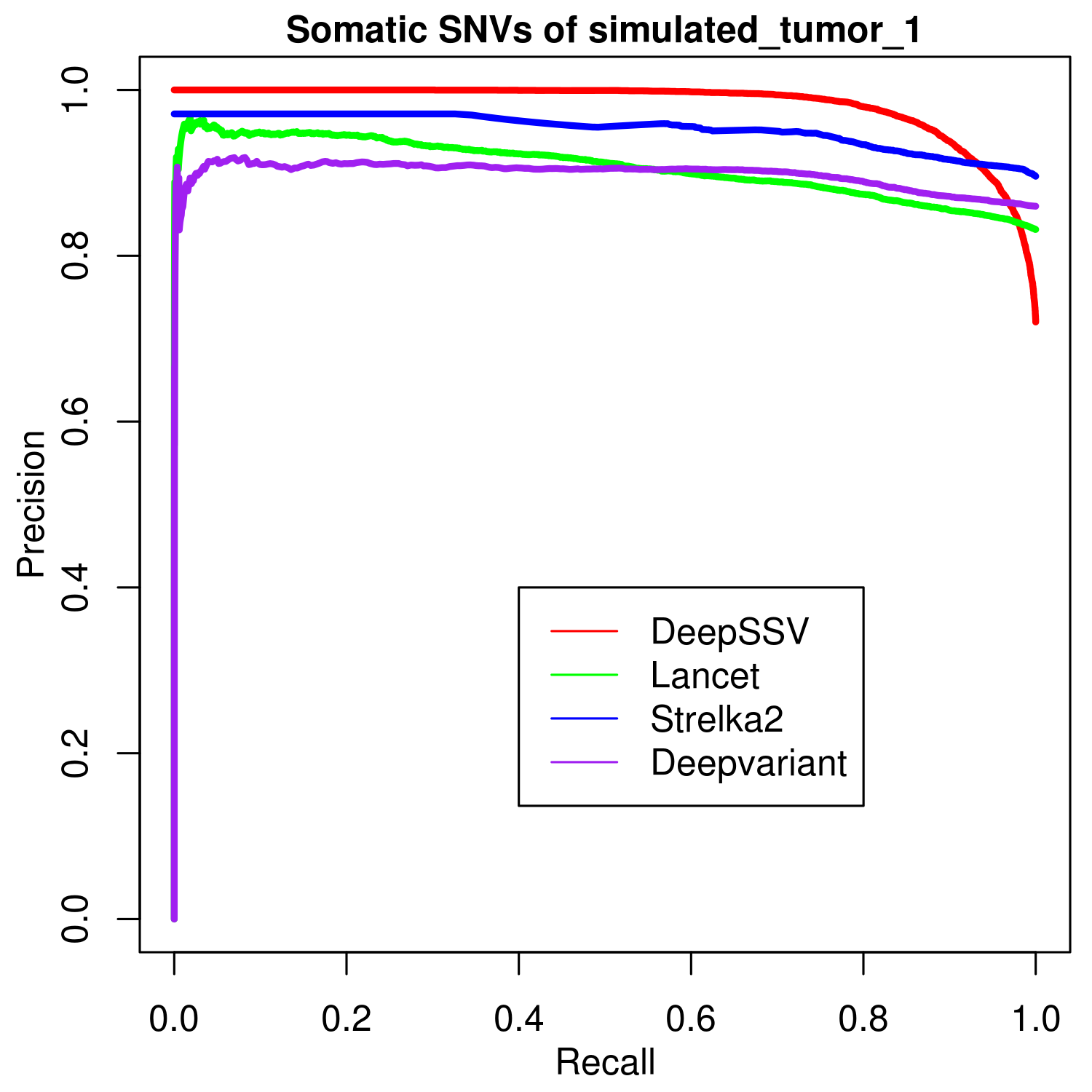

**Supplementary Figure S4.** PRROC curve of somatic callers on somatic SNVs of the simulated_tumor_1 genome.

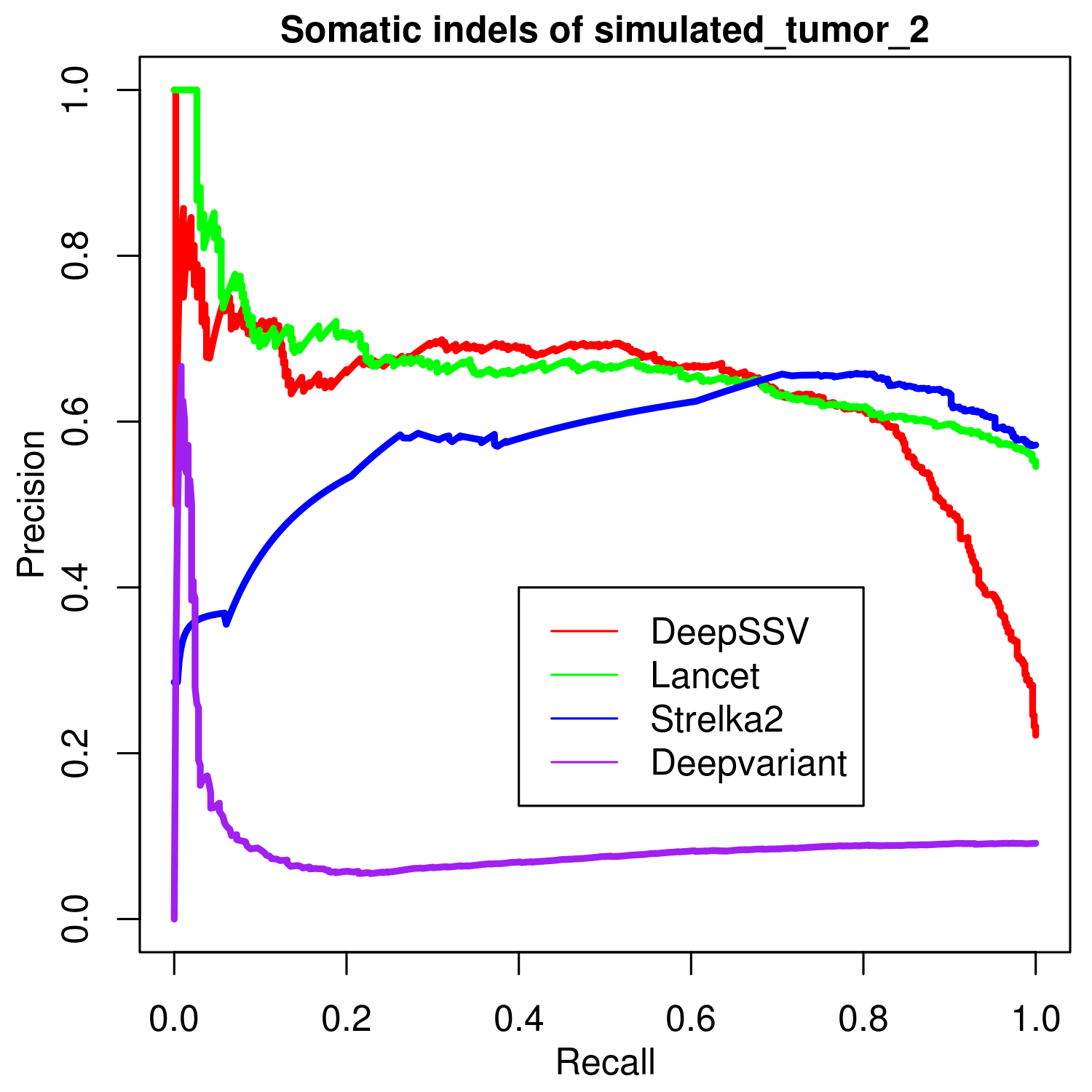

**Supplementary Figure S5.** PRROC curve of somatic callers on somatic indels of the simulated_tumor_2 genome.

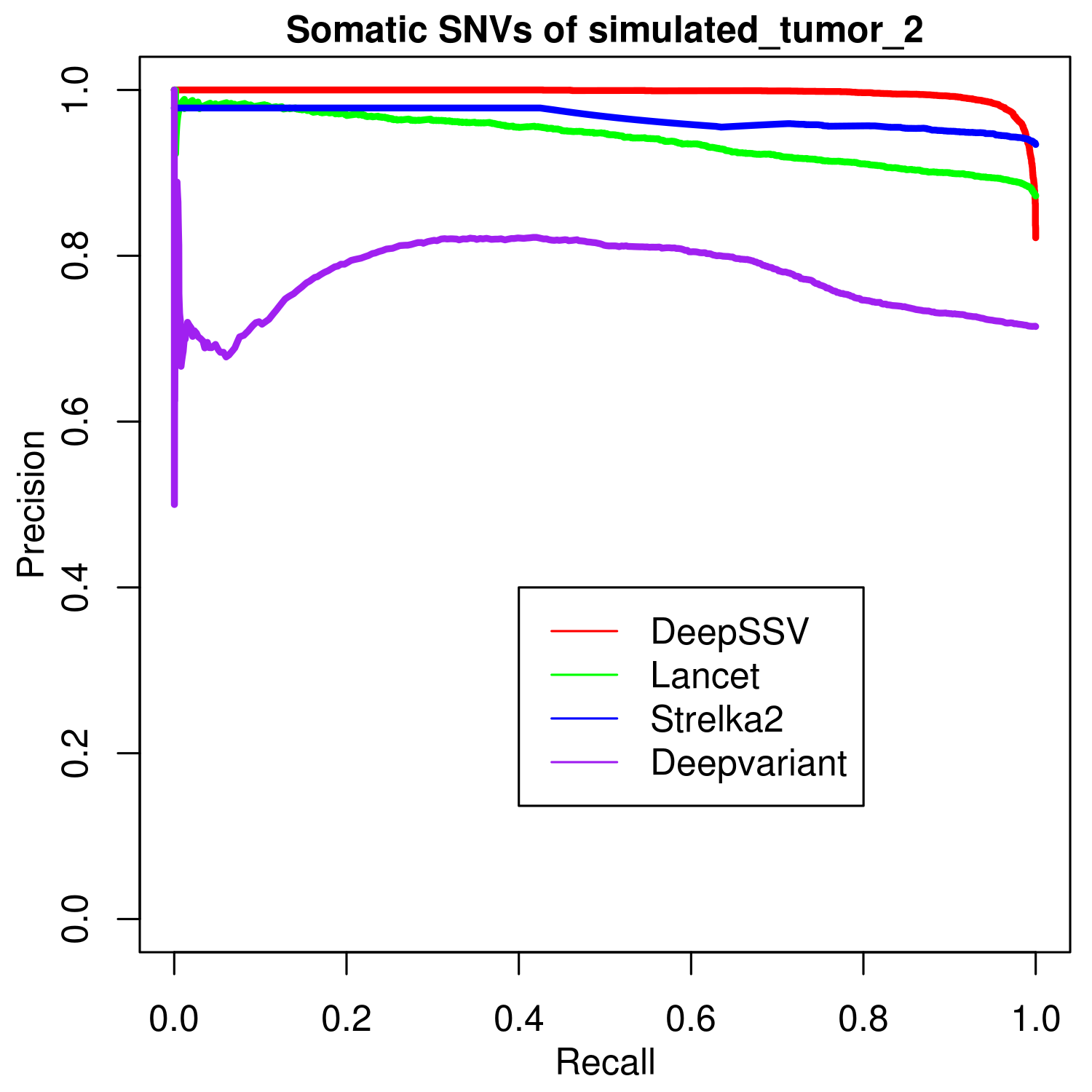

**Supplementary Figure S6.**  PRROC curve of somatic callers on somatic SNVs of the simulated_tumor_2 genome.

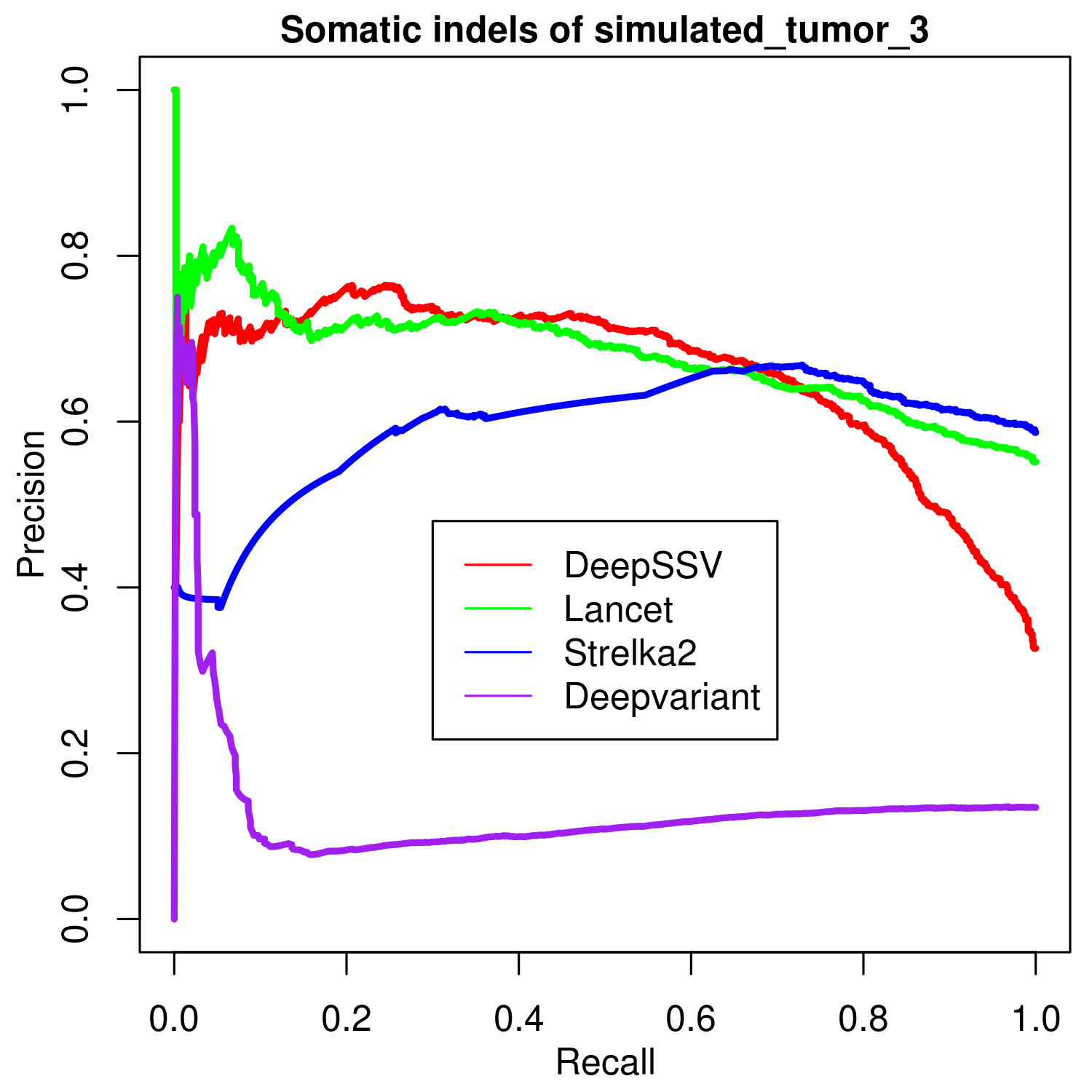

**Supplementary Figure S7.** PRROC curve of somatic callers on somatic indels of the simulated_tumor_3 genome.

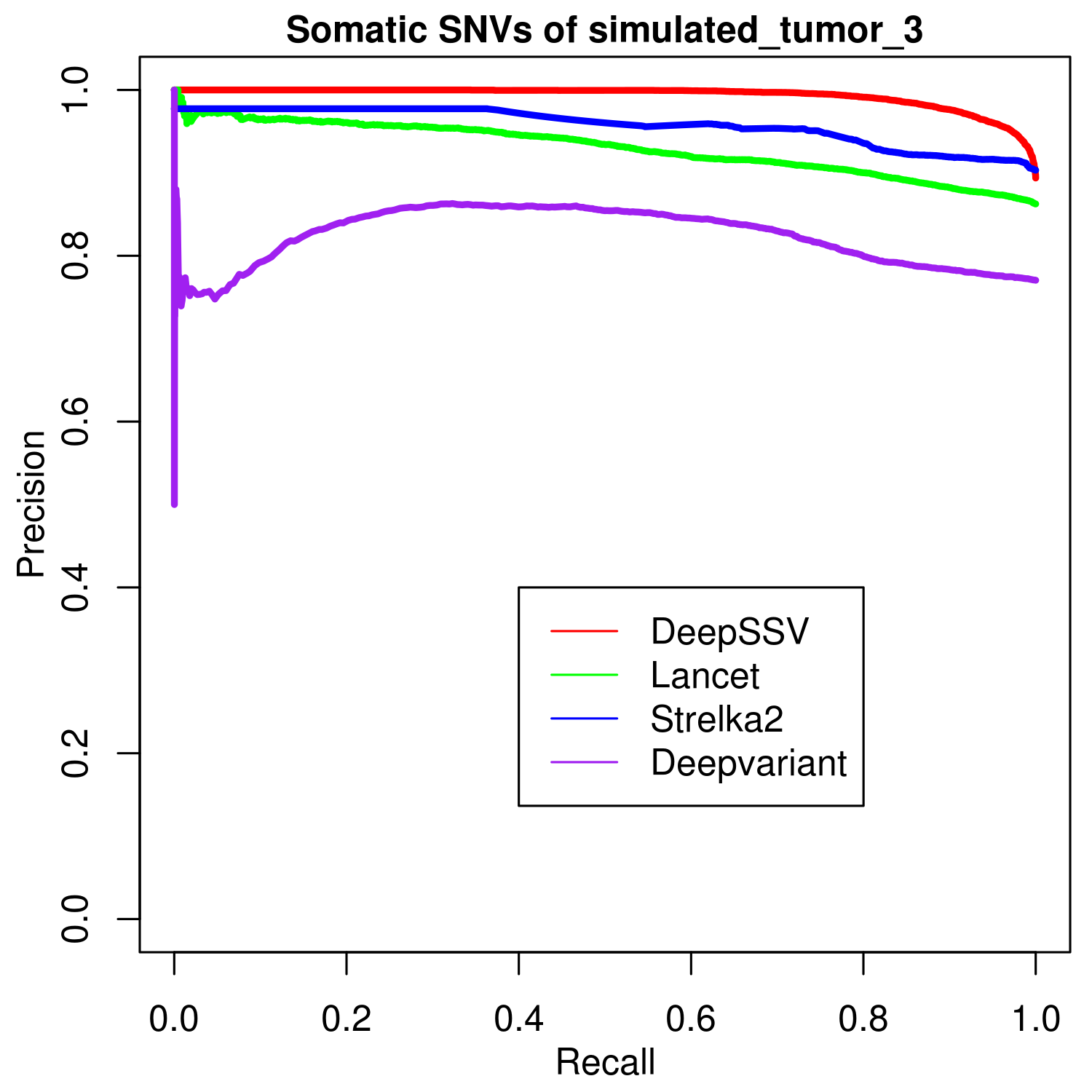

**Supplementary Figure S8.** PRROC curve of somatic callers on somatic SNVs of the simulated_tumor_3 genome.

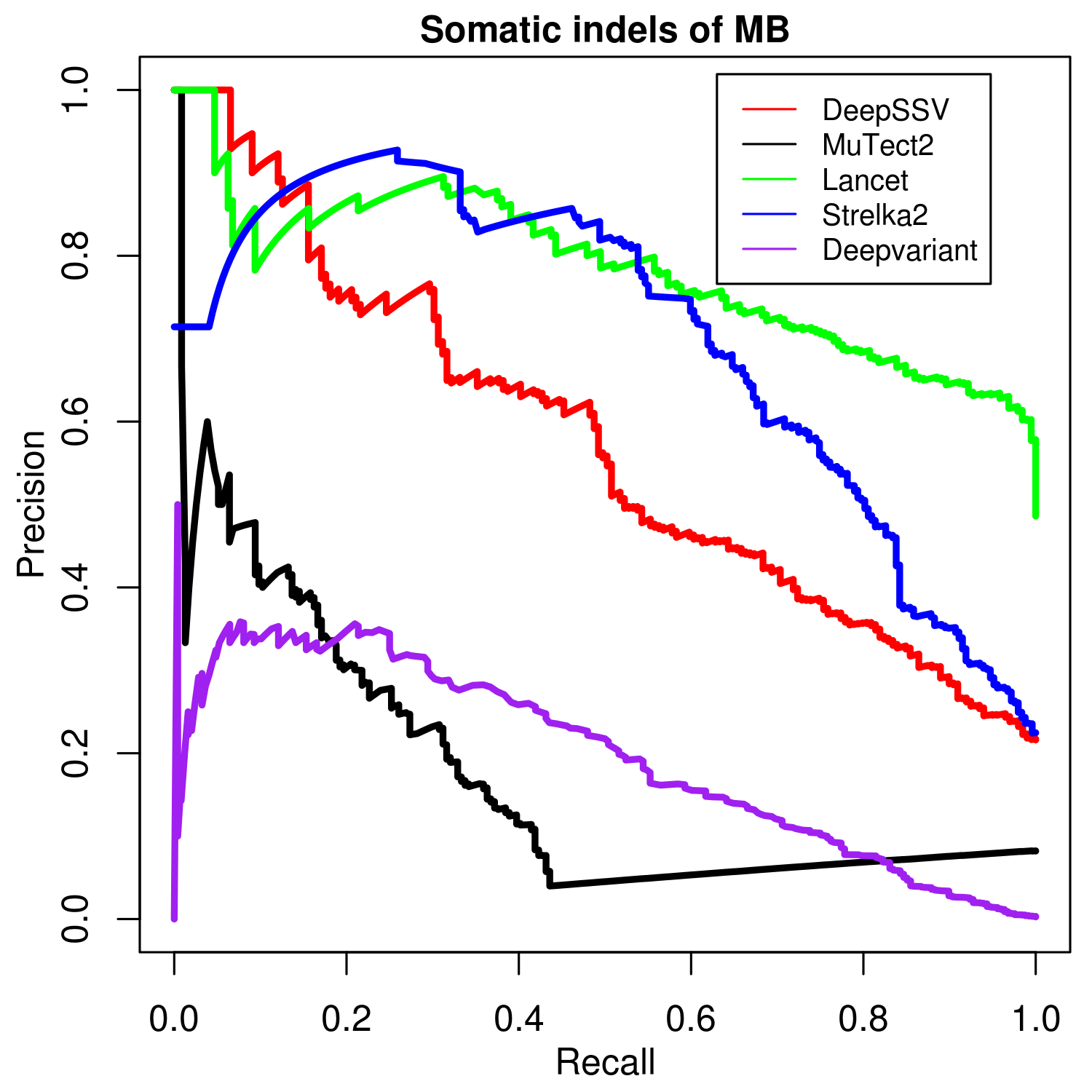

**Supplementary Figure S9.** PRROC curve of somatic callers on somatic indels of the MB genome.

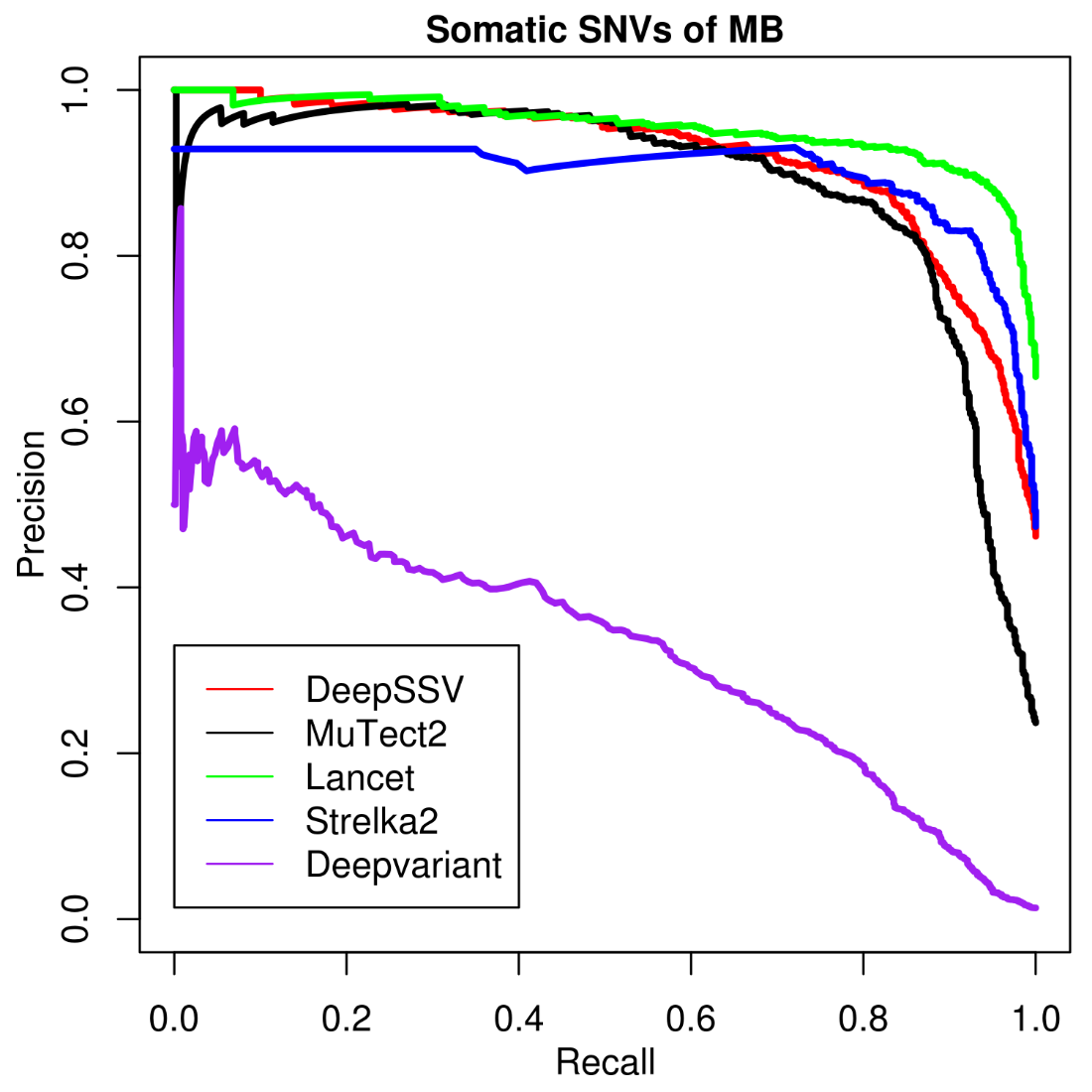

**Supplementary Figure S10.** PRROC curve of somatic callers on somatic SNVs of the MB genome.

**Supplementary Tables**

**Supplementary Table S1.** Description of pre-tumor/normal sequencing data.

| Pre-tumor (BAM) | Simulated tumor (BAM) | Sequencing machine (Illumina) | Depth | Read length | Sequencing content |
| --- | --- | --- | --- | --- | --- |
| NA12878_HiSeq1_normal | simulated_tumor_1 | HiSeq 2500 | 50 | 148 | Genome |
| NA12878_Illumina2_normal | simulated_tumor_2 | HiSeq 2000 | 50 | 101 | Genome |
| NA12878_Illumina2_normal | simulated_tumor_3 | HiSeq 2000 | 50 | 101 | Genome |

**Supplementary Table S2.** Characteristics of simulated tumor genomes.

| Simulated tumor | Mutation load | | # of sub-clones | Expected VAFs |
| --- | --- | --- | --- | --- |
|  | Somatic SNVs | Somatic indels |  |  |
| simulated_tumor_1 | 10/Mb | 1/Mb | 4 | 0.1, 0.2, 0.35 and 0.5 |
| simulated_tumor_2 | 5/Mb | 0.5/Mb | 3 | 0.2, 0.35 and 0.5 |
| simulated_tumor_3 | 10/Mb | 1/Mb | 4 | 0.1, 0.2, 0.35 and 0.5 |

10/Mb: 10 mutations per megabase.
